# Microbial and small zooplankton communities predict density of baleen whales in the southern California Current Ecosystem

**DOI:** 10.1101/2025.09.25.678658

**Authors:** Erin V. Satterthwaite, Trevor D. Ruiz, Nastassia V. Patin, Michaela N. Alksne, Len Thomas, Julie Dinasquet, Robert H. Lampe, Katherine G. Chan, Nicholas A. Patrick, Andrew E. Allen, Simone Baumann-Pickering, Brice X. Semmens

## Abstract

Understanding the density of marine mammals is important for assessing population dynamics and evaluating the impacts of human activities on these species. In this work, we examine microorganisms that may be ecologically associated, whether directly or indirectly, with baleen whales – the potential “ecological habitat” – to predict baleen whale densities via statistical relationships adjusted for seasonal patterns of occurrence. We assessed the capability of microbial and small plankton communities to predict the density of *Balaenopteridae* whales in the Southern California Current Ecosystem in each quarterly season from 2014 to 2020 using data from the California Cooperative Oceanic Fisheries Investigations (CalCOFI). Densities were estimated from visual sightings for three target species – blue, fin, and humpback whales – and microbial and small plankton communities were examined using concurrent water samples via metabarcoding of the 16S and 18S rRNA genes. We identified microbial and small plankton communities specific to each target species that were strong predictors of estimated density. Groups of 23-60 distinct Amplicon Sequence Variants (ASVs) per baleen whale species explained 81-99% of variability in estimated whale density and predicted density estimates to within ∼1 individual per 1000 *km^2^*. The predictive accuracy achieved by our approach, compared with naive seasonal carry-forward and seasonal averaging approaches to prediction, improves out-of-sample RMSE by up to an estimated 65%. Microbial and small plankton communities were characterized by 148 unique taxonomic annotations enumerated across marker genes. Of these, 20% were shared across all three species, 21% were shared by two species, and the remaining (59%) were unique to a single whale species, which suggests that there is some overlap in the ecological habitat among blue, humpback, and fin whales, but each species is also related to a distinct community of microbes and small plankton.

We also conducted a narrative review to examine existing known relationships of baleen whales with microbes or small plankton. Of the 148 unique taxonomic annotations we found to be predictive of baleen whale densities, 23% have been referenced within the existing literature exploring microbial prey, parasites, commensals, or respiratory-associated organisms of baleen whales, with matches at the genus or family level. The rest matched at higher taxonomic levels (41%) or had no documented matches in our literature search (36%). These findings suggest that some of the microbial and small plankton community predictors may be ecologically relevant, yet further studies are needed to understand how these organisms function collectively as a community and interact with whale ecology, prey, and the surrounding environment. Our results suggest that using microbial and small plankton communities to quantify the potential ecological habitat of larger organisms, like baleen whales, can enhance predictive models and may inform hypotheses about the ecological relationships between whales and the biological communities with which they co-occur.

## 1 INTRODUCTION

Large baleen whales play a vital role in marine ecosystems, providing cultural value and helping to regulate ecosystem processes (Bowen 1997, Cook *et al*. 2020). They are of particular conservation and management relevance as many populations are listed as *endangered* or *threatened* on the IUCN Red List of Threatened Species (IUCN 2025). Blue, fin, and humpback whales, among the largest and most widely distributed of the *Balaenopteridae* family, migrate to the productive coastal waters of the California Current during spring and summer months to forage on dense aggregations of krill and schooling fish (Fiedler *et al*. 1998, Fleming *et al*. 2016, Scales *et al*. 2017).

The seasonal distribution and abundance of these species in the California Current Ecosystem (CCE) is largely shaped by oceanographic conditions that influence prey availability. Blue whales preferentially consume specific krill species, favoring *Thysanoessa spinifera* over *Euphausia pacifica* (Nickels et al. 2018), and track their aggregations (Fiedler et al. 1998; Croll et al. 2005). Humpback whales are opportunistic foragers, feeding on both krill and small schooling fish such as sardine, anchovy, sand lance, and herring (Baker et al. 1985; Geraci et al. 1989; Clapham et al. 1997), with diet composition likely reflecting prey community structure and underlying oceanographic conditions (Fleming et al. 2016). Fin whales are also opportunistic foragers, though less is known about their prey preferences, seasonal distribution, and general ecology; they occur year-round in the CCE (Širovic et al. 2015) but, like blue whales, peak density occurs in summer months (Campbell et al. 2015). Across species, migratory phenology and interannual variability are closely linked to environmental conditions, with long-term acoustic studies documenting shifts in call occurrence associated with biological productivity (Oestreich et al. 2022; Ryan et al. 2025), sea surface temperature (Szesciorka et al. 2020), and broader events such as marine heatwaves and decadal-scale climate fluctuations (ZoBell et al. 2025).

The close association between baleen whales and their preferred habitat suggests that their density in the CCE may be linked to habitat-specific biological characteristics. Here, we introduce a novel approach to understanding such ecological linkages by using the community composition of microbes and small plankton to predict regional whale density, what we refer to as the “ecological habitat” (following Satterthwaite *et al*. 2023) of baleen whales.

This ecological habitat could include a suite of taxa with direct or indirect relationships to baleen whales. Direct relationships might include: prey items, parasites or commensal species that live on or in the animals; or various bacterial associates that make up the microbiome, such as those found in their gut, respiratory tract, and on their skin (Reidy *et al*. 2022; Apprill *et al*. 2010, 2014, 2017; Guass *et al*. 2016; Herwig & Staley 1986; Domínguez-Sánchez *et al*. 2022; Glaeser *et al*. 2022; Vendl *et al*. 2020, Measures 1993; Rausch & Rice 1970; Jo *et al*. 2017; Kane *et al*. 2008; Félix *et al*. 2006; Ten *et al*. 2022). Whale microbiomes have been found to significantly differ from the microbial community in the surrounding seawater (Vendl *et al*. 2020, Apprill *et al*. 2017). For example, distinct bacterial taxa were associated with the skin of humpback whales across the North Pacific (Apprill *et al*. 2010). Additional studies have observed parasites, viruses, and epibiotic fauna specific to whales (Measures 1993; Rausch & Rice 1970; Jo *et al*. 2017; Kane *et al*. 2008; Félix *et al*. 2006; Ten *et al*. 2022; Domínguez-Sánchez *et al*. 2022; Glaeser *et al*. 2021). There also may be taxa that are part of the “ecological habitat” and indirectly associated with baleen whales, such as organisms at the base of the food web (e.g., prey of whale prey), parasites, commensals of organisms associated with whales, or other organisms that co-occur with whales due to coinciding seasonal or oceanographic affiliations.

Monitoring the distributions of baleen whales over time is necessary for tracking population changes and understanding the effects of human activity on these species, both of which are important for effective conservation and management. Baleen whales are monitored using a combination of visual (Barlow and Forney, 2007; Campbell *et al*. 2015) and acoustic (Širović *et al*. 2015) surveys, satellite imagery (Cubaynes *et al*. 2019), tags (Irvine *et al*. 2014), and, increasingly, genetic methods (Baker *et al*. 2018). Sampling whales and other marine mammals is challenging due to their wide-ranging and often patchy distributions, low encounter rates, brief or intermittent surface cues, deep-diving or cryptic behaviors, and the high logistical, financial, and permitting costs of direct or invasive sampling methods.

Because large baleen whales are difficult to sample directly, yet are closely linked to their habitat and associated biota, we investigated whether microbial and small plankton communities could serve as proxies for predicting their density in the southern California Current Ecosystem. Understanding the ecological habitat of baleen whales in a holistic, community-based way has been challenging due to variety and the microscopic size of many associated organisms, as well as the contemporaneous sampling needed to assess many different types and sizes of organisms.

Here, we leverage eDNA metabarcoding to characterize baleen whale ecological habitat and to predict their seasonal density in the California Current Ecosystem. We used two genetic markers, the 16S and 18S ribosomal RNA genes, to capture a wide range of prokaryotic and eukaryotic microbes as well as small metazoan zooplankton like copepods. We use the terms “communities of microbes and small zooplankton” or “microbial and small plankton communities” to broadly encompass the ecological communities of small organisms associated with of baleen whales such as bacteria, archaea, single-celled phytoplankton and small, metazoan zooplankton. We focus our analyses on blue, fin, and humpback whales, given that they are highly abundant in the California Current Ecosystem, forage at low trophic levels, and have been shown to have existing connections to various microbes. Additionally, concurrent eDNA samples and visual sightings of baleen whales exist from the California Cooperative Oceanic Fisheries Investigations (CalCOFI), the longest integrated marine ecosystem observing program in the world.

## 2 METHODS

### 2.1 SAMPLING AREA

The Southern California Bight region is situated in the southern portion of the California Current Ecosystem (CCE), a productive upwelling system that supports important baleen whale species. The California Cooperative Oceanic Fisheries Investigations (CalCOFI), one of the longest running integrated marine ecosystem monitoring programs in the world, has systematically sampled the physics, chemistry, and biology of the CCE since 1949.

Quarterly CalCOFI sampling consists of 75 stations along six “core area” transects that extend from San Diego, CA to north of Point Conception (Morro Bay, CA) and include coastal stations (∼50 m depth) and stations within the core of the California Current (CC) out to ∼250 to 550 km offshore. The transects are spaced approximately 40 nautical miles (nm) apart. Along the transect lines, stations are spaced approximately 40 nm apart for offshore stations and 20 nm apart for coastal stations. In this project we utilize data collected from the core sampling area comprising stations located on CalCOFI lines 93.3 to 76.7 (SE corner: 32.956, -117.305; NE corner: 35.088, -120.777; NW corner: 33.388, -124.323; SW corner: 29.846, -123.587) between 2014 and 2020.

### 2.2 ENVIRONMENTAL DNA COLLECTION AND AMPLICON SEQUENCING

From select stations on CalCOFI cruises, 743 DNA samples were collected within the core sampling area from 2014-2020 as part of the NOAA-CalCOFI Ocean Genomics (NCOG) time series (James et al. 2022). We sampled key stations on lines 80 and 90, as well as basin stations, to provide an onshore-offshore gradient across two transects. Seawater (mean = 3.3 L) was collected from the near-surface (normally 10 m) and the subsurface chlorophyll maximum layer and filtered onto 0.22 µm Sterivex^TM^ filters that were immediately flash frozen in liquid nitrogen and stored at -80℃. DNA was extracted with the Macherey-Nagel NucleoMag Plant kit on an Eppendorf epMotion 5075TMX and assessed on a 1.8% agarose gel.

Amplicon libraries separately targeting the V4-V5 region of the 16S rRNA gene and both the V4 and V9 regions of the 18S rRNA genes were constructed via a one-step PCR with the TruFi DNA Polymerase PCR kit. For 16S, the 515F-Y (5′-GTG YCA GCM GCC GCG GTA A-3′) and 926R (5′-CCG YCA ATT YMT TTR AGT TT-3′) primer set was used (Parada *et al*. 2016). For 18S-V4, the V4F (5′-CCA GCA SCY GCG GTA ATT CC-3′) and V4RB (5′-ACT TTC GTT CTT GAT YR-3′) primer set modified from Berdjeb *et al*. (2018) was used. For 18S-V9, the 1389F (5′-TTG TAC ACA CCG CCC-3′) and 1510R (5′-CCT TCY GCA GGT TCA CCT AC-5′) primer set was used (Amaral-Zettler *et al*. 2009).

Each reaction was performed with an initial denaturing step at 95°C for 1 minute followed by 30 cycles of 95°C for 15 seconds, 56°C for 15 seconds, and 72°C for 30 seconds. 2.5 µL of each PCR reaction was run on a 1.8% agarose gel to confirm amplification, then PCR products were purified with Beckman Coulter AMPure XP beads following the manufacturer’s instructions. PCR quantification was performed in duplicate using the Invitrogen Quant-iT PicoGreen dsDNA Assay kit. Samples were then combined in equal proportions into multiple pools followed by another 0.8x AMPure XP bead purification on the final pool. DNA quality of each pool was evaluated on an Agilent 2200 TapeStation, and quantification was performed with the Qubit HS dsDNA kit. Each 16S or 18S pool was sequenced on an Illumina MiSeq (2 x 300 bp for 16S and V4 or 2 x 150 bp for V9) except for the one pool for the 2014-2016 euphotic zone V9 samples, which was run on an Illumina NextSeq 500 (Mid Output, 2 x 150 bp).

Amplicons were analyzed with QIIME2 v2019.10 (Bolyen *et al*. 2019). Briefly, paired-end reads were trimmed to remove adapter and primer sequences with cutadapt (Martin 2011). Trimmed reads were then denoised with DADA2 to produce amplicon sequence variants (ASVs). Each MiSeq run was denoised with DADA2 separately to account for different error profiles in each run then merged. Taxonomic annotation of ASVs was performed with the q2-feature-classifier naïve bayes classifier using the SILVA database (Release 138) for 16S ASVs and the PR^2^ database (v4.13.0) for 18S ASVs (Bokulich *et al*. 2018; Guillou *et al*. 2012; Pedregosa *et al*. 2011; Pruesse *et al*. 2007).

### 2.3 VISUAL SURVEYS OF BALEEN WHALES

Since 2004, visual sightings of baleen whales have been recorded during cruises along CalCOFI transects. In this project, we focus on data from 2014 to 2020 collected contemporaneously with the NCOG data described above in Section 2.2. The dataset includes marine mammal monitoring effort from 25 individual CalCOFI cruises spanning all four seasons and limited to the core sampling area.

Visual monitoring effort was conducted in “passing mode” and adapted from standard line-transect marine mammal survey protocols (Barlow 1997, Barlow and Forney 2007) following methods outlined in Campbell *et al*. (2015). Two trained marine mammal observers used 7x50 Fujinon binoculars to observe and record marine mammals during daylight hours as the ship transited between CalCOFI stations. Observers systematically recorded species identification, group size estimates, reticle position below the horizon, angle relative to the bow, latitude and longitude, ship’s heading, sea state, swell height and visibility. Survey effort was suspended when sea state was greater than Beaufort 6 or when visibility less than 1 km.

In this project, whale sightings were only included when classified as both “on-effort” and “on-transect”. The on-effort criteria was met when two observers were actively scanning while the vessel was traveling above 10 knots in a sea state below Beaufort 6 with greater than or equal to 1 km visibility. The on-transect criteria was met when sightings were along one of the core CalCOFI transect lines. “Off-transect” sightings (which were not used here) occurred during north/south coastal transits or between transect lines, during transits to and from port, and during any other non-standard transit.

### 2.4 ESTIMATED BALEEN WHALE DENSITIES

The marine mammal visual survey data was used to estimate density (number of individuals per 1000 km^2^) using multiple covariate distance sampling methods (Marques *et al*. 2007). This analysis involved two stages: (1) estimating each species’ detectability as a function of factors potentially affecting sighting conditions; (2) estimating species’ density per cruise given the number cited and the estimated detectability. The major advantage of this approach over simply using sighting rates (number of individuals detected per unit survey effort) is that it can account for differences in sighting rates that are caused by differences in detectability (for example, if sighting conditions tend to be worse in winter) that might otherwise confound downstream inferences about the relationship between whales and their ecological habitat. Distance sampling analyses were undertaken using the Distance R package (Miller *et al*. 2019).

Distance sampling methods use the distribution of perpendicular distances of observed animals to estimate a “detection function” (i.e., probability of detection as a function of perpendicular distance and other covariates), and from this the average probability of detection within the surveyed strips (Buckland *et al*. 2001). For each sighting, the measured reticle position and angle relative to bow were used, together with knowledge of observer eye height above the water, to calculate perpendicular distance for each sighting. Following Campbell *et al*. (2015), a perpendicular truncation distance of 2400 m was used. For each species, candidate detection functions were fitted incrementally using forward selection, starting with a key function and adding terms if the resulting model had a lower Akaike Information Criterion (AIC) score. Key functions were half-normal and hazard rate. In one set of analyses the terms added were cosine (with half-normal) or polynomial (with hazard rate) adjustment terms. In another set of analyses, the terms added were covariates affecting the scale parameter of the key functions: Beaufort sea state, swell height, observer height above water (as a factor with 3 levels) and group size. The final model chosen for inference was the one with lowest AIC over both sets. A list of all candidate models is given in supplemental materials. Goodness of fit was assessed visually by comparing the fitted detection function with histograms of observed distances, and using a Cramér–von Mises test.

Given a fitted detection function, estimated density per cruise, denoted 𝑦_𝑖_, for each species was calculated as:

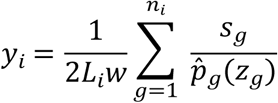

where 𝐿_𝑖_is the transect length on cruise *i*, *w* is the truncation distance, 𝑛_𝑖_is the number of detections of the species on survey *i*, 𝑠_𝑔_ is the group size of the *g*th detection, and *p*^_𝑔_(𝑧_𝑔_) is the estimated detection probability of the *g*th detection given its covariates 𝑧_𝑔_. This detection probability is computed by averaging the detection function over the perpendicular distances:

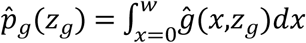

where ĝ(𝑥,𝑧_𝑔_) is the estimated detection function and *x* is perpendicular distance. Variance in estimated density was calculated by combining variance in the estimated detection function and variance in detection rate between transect lines, as detailed by Marques and Buckland (2004).

### 2.5 ANALYSIS OF AMPLICON RELATIVE ABUNDANCES

We removed rare ASVs present in under 1% of samples across all cruises and abundant ASVs present in more than 99% of samples across all cruises. Under the assumption that all remaining ASVs were physically present across samples and cruises, Geometric Bayesian multiplicative count zero imputation (Martín-Fernández *et al*. 2015) was used to estimate relative abundances for non-detections.

To match the spatial resolution of the whale density estimates, amplicon relative abundances were aggregated to the cruise level by weighted (geometric) averaging. In detail, if 𝑥_𝑖𝑗𝑘𝑙𝑚_denotes the relative abundance of ASV 𝑗 from the sample taken at station 𝑙 on transect 𝑘 and depth 𝑚 on cruise 𝑖, relative abundances were aggregated across depth and sampling location by taking a weighted geometric mean:

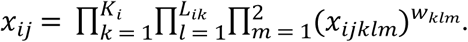

Weights 𝑤_𝑘𝑙𝑚_were inversely proportional to spatial sampling density with respect to geolocation and maximized 𝛼-diversity with respect to depth; the latter criterion resulted in weights slightly favoring samples taken at max chlorophyll-*a* depth. The resulting quantity measures the average relative abundance of ASV 𝑗 observed across samples collected on cruise 𝑖.

The centered log-ratio (CLR) transformation (Aitchison 1982) was then applied to average relative abundances 𝑥_𝑖𝑗_. The “typical” average relative abundance taken across ASVs on cruise 𝑖 is given by the geometric mean:

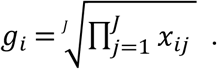

The CLR transformation is defined as the natural logarithm of the ratio:

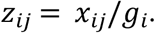

This captures the factor by which the average relative abundance of a particular ASV deviates from the typical average relative abundance across all ASVs on a given cruise; for example, value of 𝑧_𝑖𝑗_ = 2 indicates that ASV 𝑗 is twice as abundant as the typical ASV on cruise 𝑖.

The above aggregations and transformations were performed separately for the 16S, 18S-V4, and 18S-V9 markers, yielding three sets of 𝑧_𝑖𝑗_; this level of detail is omitted from the notation.

### 2.6 SEASONAL LOGRATIOS

Seasonality was removed from the whale density and amplicon data with a secondary logratio transformation using the respective seasonal averages. The seasonal geometric means, written as functions of the observation index (cruise) 𝑖, are:

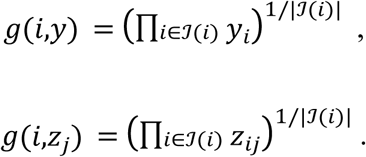

In these expressions ℐ(𝑖) is an index set comprising the indices of all observations made in the same quarter as observation 𝑖. The seasonally-adjusted estimated whale density and the seasonally-adjusted average amplicon relative abundance are then 𝑦_𝑖_/𝑔(𝑖,𝑦) and 𝑧_𝑖𝑗_/𝑔(𝑖,𝑧_𝑗_), respectively. The resulting quantities are best interpreted as deviations from seasonal averages; for example, a value of 𝑦_1_/𝑔(1,𝑦) = 0.5 would indicate that on the first cruise, observed density was half of the seasonal average for the corresponding quarter.

### 2.7 LOG-CONTRAST MODEL FRAMEWORK

We formulated a log-contrast-type model (Aitchison and Bacon-Shone, 1984) to identify and estimate statistical relationships. This model expresses the seasonally-adjusted estimated densities as linear functions of the seasonally-adjusted average amplicon relative abundances:

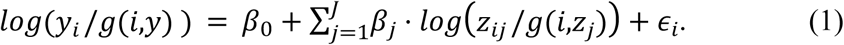

Due to the logratio transformations, the coefficients capture multiplicative changes in median estimated density associated with multiplicative changes in relative abundances, after adjusting for seasonality and assuming error normality. For example, a twofold change in the relative abundance of ASV 𝑗, relative to its seasonal average, is associated with a change in median density, relative to its seasonal average, of a factor of 2^𝛽𝑗^. We specified separate models for each whale species of interest and each marker, amounting to 9 models in total.

### 2.8 VARIABLE SELECTION AND PARAMETER ESTIMATION

A partial least squares (PLS) latent variable framework (Wold *et al*. 2001) was used for variable selection and parameter estimation. PLS allows for estimation of the full set of model coefficients even when least squares is ill-posed due to the number of covariates (ASVs) exceeding the number of samples (cruises); it is the only available framework in this setting that does not require selecting a subset of ASVs smaller than the number of cruises (25) in order to fit the model. Writing the log-contrast model (Eqn. 1) in linear model form 𝑌 = 𝑍𝛽 + 𝜖, the PLS framework stipulates a set of latent variables or “components”:

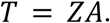

The columns 𝑎_𝑘_ of 𝐴 or component “loadings” are estimated sequentially by component 𝑘 = 1, …, 𝐾 by maximizing the correlation with the response and the variance of the latent component, subject to an orthogonality constraint with respect to previous latent components and a unit-norm constraint (when 𝑘 = 1 the orthogonality constraint is omitted):

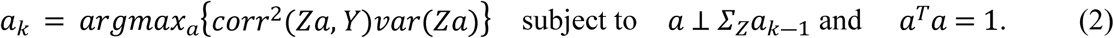

The SIMPLS algorithm (De Jong 1993) was used to compute estimates of the loading matrix 𝐴. Subsequently, a linear model was fit with the latent components 𝑇 as covariates and least squares estimates were back-propagated to obtain coefficient estimates for the model as originally specified:

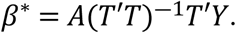

The log-contrast models (Eqn. 1) were specified using a small subset of candidate amplicons identified through variable selection to improve model interpretability and prioritize identification of relatively stronger associations. The selection procedure resulted from applying the stability selection method of Meinshausen and Bühlmann (2010) to sparse partial least squares (sPLS) estimates of 𝐴 (Chun & Keleş 2010). The sPLS method introduces an 𝐿_1_ penalty to the PLS optimization problem (Eqn. 2), which has the effect of shrinking small component loadings 𝑎_𝑘𝑗_ to exactly zero and inducing sparsity in the loading matrix 𝐴. Chun & Keleş (2010) approximate the solution to the resulting problem using a surrogate approach with a hyperparameter 𝜆 controlling the degree of sparsity in the sPLS estimate; this results in a sparse coefficient estimate 𝛽^𝜆^. Stability selection is a computationally-intensive procedure that traverses the problem of hyperparameter tuning by estimating the selection probability of each variable for a “path” of hyperparameter values in a specified region 𝛬. Selection probability estimates are obtained by computing 𝛽^𝜆^repeatedly from subsamples of the data.

We used leave-one-out partitions to estimate, for each hyperparameter 𝜆 ∈ 𝛬 and each candidate amplicon 𝑗, the probability of selecting that amplicon using the sPLS method:

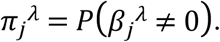

The “stable set” consists of frequently-selected amplicons, specifically those variables whose estimated selection probability exceeds 𝜋_𝑚𝑎𝑥_ for at least one 𝜆 ∈ 𝛬:

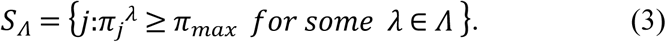

Meinshausen and Bühlmann (2010) provide heuristics for choosing the region 𝛬 to control the expected number of falsely selected variables (per-family error rate); we determined 𝛬 according to their method in order to control the per-family error rate at 0.5.

In the context of our model, the sPLS coefficient estimate 𝛽^𝜆^ depends not only on the sparsity hyperparameter 𝜆 but also on the number 𝐾 of latent components. We addressed this by estimating stable sets 𝑆_𝛬_^𝐾^for 𝐾 = 4, …, 12 and choosing the number of components that optimized mean square prediction error estimated from leave-one-out partitions of the data. We re-computed seasonal adjustments when forming data partitions so that subsamples did not incorporate information about held-out observations via seasonal averages. Once the stable set 𝑆 was estimated for each model, we computed SIMPLS estimates of the model coefficients with the number of latent components 𝐾 used to determine the stable set.

### 2.9 MODEL VALIDATION

We sought to further assess the consistency of the variable selection procedure by comparing stable sets obtained under perturbations of the data partitions used to estimate selection probabilities. An “outer validation” was performed by constructing a set of nested leave-one-out partitions and performing the entire stability selection procedure holding out one observation (cruise) at a time. This yielded stable sets 𝑆_1_, … , 𝑆_𝑛_ (one from holding out each of the 𝑛 = 25 cruises) which we then compared for consistency using a thresholded Jaccard index:

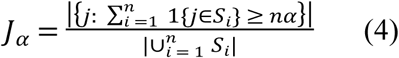

This measures the proportion of amplicons selected at least 𝛼% of the time among the stable sets obtained from the outer validation procedure. We chose 𝛼 = 0.5 to look at the proportion of amplicons selected more often than not across validation runs as a measure of consistency of the variable selection procedure.

### 2.10 NARRATIVE REVIEW OF DIRECT RELATIONSHIPS BETWEEN BALEEN WHALES AND MICROBES/SMALL ZOOPLANKTON

A narrative review was conducted to identify known direct relationships between blue, humpback, and fin whales and bacteria, microbes, and small plankton from the existing literature for comparison with potential relationships identified by our models. The review focused on studies that examine microbial and planktonic interactions with these baleen whale species. The Publish or Perish software was utilized to search the academic literature database Google Scholar for peer-reviewed articles and reports. Specific title-keyword combinations were searched to capture relevant studies related to bacteria, microbes, and plankton associated with blue, humpback, and fin whales. Search terms included: “humpback whale” in the title with “bacteria” or “plankton” as keywords; “blue whale” in the title with “bacteria” or “plankton” as keywords; and “fin whale” in the title with “bacteria” or “plankton” as keywords.

The titles of these papers were reviewed to assess relevance to the research question, focusing on studies that explored direct connections between baleen whales and microbes, bacteria, or plankton. Studies that examined direct interactions between microbes or small plankton and baleen whales were selected. Interactions included topics such as: baleen whale health and disease, including microbial diversity and baleen whale pathology; feeding ecology, including studies that analyzed prey-plankton dynamics and their relationship with baleen whale foraging behavior; baleen whale microbiome, such as studies on respiratory, gut, or skin microbiomes; baleen whale parasitology and pathology, which examined diseases and parasites associated with baleen whale health; and baleen whale strandings and carcasses. Studies unrelated to direct baleen whale-microbe relationships were excluded from the review. This search strategy yielded 18 relevant papers.

These selected papers were examined for relevant information that was entered into a structured table that contained the following information: citation, title of the paper, url/link to the paper, notes, whale species (e.g., blue, humpback, or fin whales), type of relationship between the whale species and bacteria, microbes, or plankton, type of sample or data used in the study, location, method, taxonomic classification of the bacteria/microbe/small plankton (Kingdom/Domain, Phylum, Class, Order, Infraorder, Family, Genus, Accepted Name).

For each microbial taxon mentioned in the selected studies, a detailed taxonomic classification was retrieved programmatically in Python using the World Register of Marine Species (WoRMS) API and programmatically in R using the NCBI (National Center for Biotechnology Information) database (Supplementary table 4b).

## 3 RESULTS

Qualitative assessment of microbial community data, baleen whale sightings, and density estimates identified spatial and temporal patterns in both baleen whale sighting rates and density estimates across species, and relatively homogeneous sampling across space and time suggests minimal bias in measurement of microbial communities (Section 3.1). Statistical models relating baleen whale density estimates to microbial community data identified small sets of ASVs that explained a large proportion of variation in whale density estimates after adjusting for seasonality (Section 3.2). The relative abundances of taxa within these subcommunities provide accurate predictions of density of target baleen whale species, and community predictors outperform naive forecasting methods (Section 3.3). Through a narrative review of existing literature, we found some of the taxa in our study have also been previously documented as microbial associates of baleen whales (Section 3.4). Lastly, despite some taxonomic overlap in microbial subcommunity predictors of blue, humpback, and fin whales, our models suggest that each species is related to a distinct subcommunity (Section 3.5).

### 3.1 DATASETS

Figure 1 shows the locations along CalCOFI survey transects of (a) NCOG samples sequenced and (b) on-effort whale sightings used in the analysis; sightings tended to occur nearer to shore. The NCOG samples were distributed fairly uniformly across space and time (Table 1); data for a given cruise and genetic marker typically comprised 20-40 samples collected across 5-6 transects during a 2-3 week period. However, there is some variation in the number of samples sequenced by cruise – with as few as 14 samples collected in spring 2014 and winter 2019 and as many as 56 collected in winter 2018 – as well as in the length of the survey period and the spatial distribution of sampling locations.

**Figure 1.**
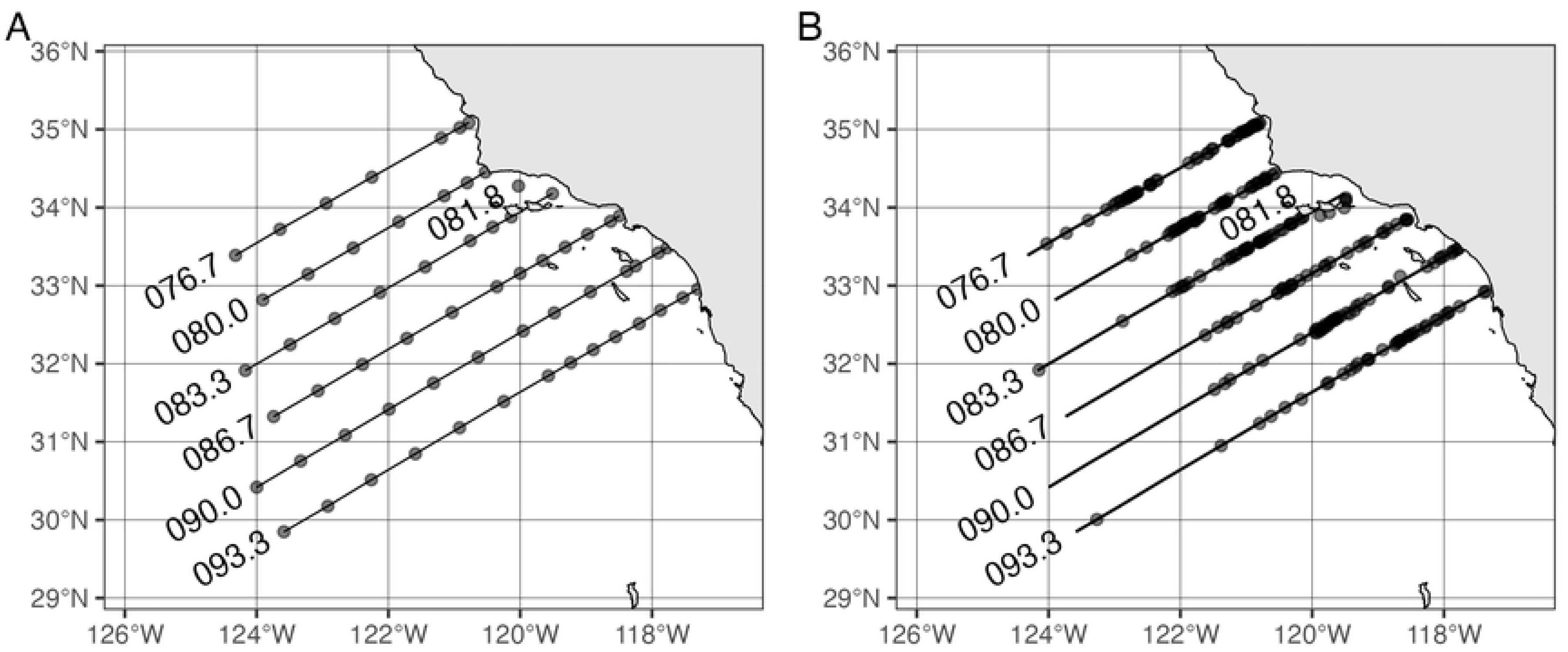
(A) Sampling locations for NCOG samples used in the analysis and (B) sighting locations recorded from visual survey data for target species during 2014-2020. All on-effort, on-transect sightings are shown; in the line transect analysis, the small number of sightings at perpendicular distances greater than 2400 m were truncated.

**Table 1.** Counts of NCOG samples and CalCOFI transects per cruise by marker.

| Year | Season | Sampling start date | Sampling end date | Samples (16S) | Transects (16S) | Samples (18SV4) | Transects (18SV4) | Samples (18SV9) | Transects (18SV9) |
| --- | --- | --- | --- | --- | --- | --- | --- | --- | --- |
| 2014 | winter | 2014-01-29 | 2014-02-04 | 14 | 2 | 12 | 2 | 12 | 2 |
| 2014 | spring | 2014-04-02 | 2014-04-16 | 32 | 6 | 32 | 6 | 32 | 6 |
| 2014 | summer | 2014-07-06 | 2014-07-19 | 26 | 4 | 26 | 4 | 24 | 4 |
| 2014 | fall | 2014-11-08 | 2014-11-22 | 26 | 6 | 24 | 6 | 24 | 6 |
| 2015 | winter | 2015-01-15 | 2015-01-29 | 28 | 7 | 28 | 7 | 28 | 7 |
| 2015 | spring | 2015-04-08 | 2015-04-17 | 14 | 2 | 14 | 2 | 14 | 2 |
| 2015 | summer | 2015-07-08 | 2015-07-23 | 24 | 6 | 22 | 6 | 24 | 6 |
| 2015 | fall | 2015-11-01 | 2015-11-10 | 22 | 5 | 22 | 5 | 22 | 5 |
| 2016 | winter | 2016-01-09 | 2016-01-20 | 22 | 6 | 22 | 6 | 22 | 6 |
| 2016 | spring | 2016-04-01 | 2016-04-14 | 22 | 6 | 22 | 6 | 22 | 6 |
| 2016 | summer | 2016-07-10 | 2016-07-24 | 18 | 5 | 18 | 5 | 18 | 5 |
| 2016 | fall | 2016-11-06 | 2016-11-20 | 26 | 6 | 26 | 6 | 26 | 6 |
| 2017 | winter | 2017-01-06 | 2017-01-18 | 38 | 7 | 32 | 6 | 38 | 7 |
| 2017 | spring | 2017-03-29 | 2017-04-12 | 32 | 6 | 32 | 6 | 32 | 6 |
| 2017 | summer | 2017-08-02 | 2017-08-13 | 34 | 5 | 26 | 4 | 34 | 5 |
| 2017 | fall | 2017-11-09 | 2017-11-22 | 32 | 5 | 20 | 4 | 32 | 5 |
| 2018 | winter | 2018-02-01 | 2018-02-09 | 56 | 6 | 56 | 6 | 56 | 6 |
| 2018 | spring | 2018-04-06 | 2018-04-20 | 32 | 6 | 32 | 6 | 30 | 6 |
| 2018 | summer | 2018-06-11 | 2018-06-23 | 32 | 7 | 32 | 7 | 32 | 7 |
| 2018 | fall | 2018-10-14 | 2018-10-27 | 34 | 6 | 34 | 6 | 34 | 6 |
| 2019 | winter | 2019-02-07 | 2019-02-12 | 14 | 4 | 14 | 4 | 14 | 4 |
| 2019 | spring | 2019-04-03 | 2019-04-17 | 32 | 7 | 32 | 7 | 32 | 7 |
| 2019 | summer | 2019-07-11 | 2019-07-26 | 44 | 7 | 44 | 7 | 44 | 7 |
| 2019 | fall | 2019-11-05 | 2019-11-17 | 34 | 7 | 36 | 7 | 36 | 7 |
| 2020 | winter | 2020-01-05 | 2020-01-18 | 26 | 6 | 26 | 6 | 26 | 6 |
| 2020 | spring | 2020-07-14 | 2020-07-25 | 26 | 6 | 26 | 6 | 26 | 6 |
| 2020 | summer | 2020-10-12 | 2020-10-22 | 32 | 6 | 32 | 6 | 32 | 6 |

*Counts reflect the samples used in the analysis rather than the total number of samples collected during the survey;* *similarly, start and end dates indicate the earliest and latest dates of samples included in the analysis.*

Supplementary Table 1 shows NCOG sample counts by cruise and transect, providing a more fine-grained look at the spatial distribution of these samples. Certain transects – namely, 080.0, 090.0, and 093.3 – were consistently sampled more densely. Supplementary tables 2a-c list the unique ASVs retained in the analysis after filtering out rare and ubiquitous ASVs for the 16S, 18S-V4, and 18S-V9 markers, along with their taxonomic classifications, if known. There were 6234, 6824, and 9511 such “candidates”, respectively.

Whale sightings exhibited clear seasonal variation (Table 2): blue whales were the most seasonal and rarely sighted except in summer; fin whales were also more common in summer, but sighted throughout the year in most years; humpback whales were sighted year-round but in greater numbers during spring. Sample sizes for the distance sampling analysis, and tables showing AIC values for fitted detection function models are given in Supplementary tables 5a-b. The selected (i.e., lowest-AIC) models were half-normal with one cosine adjustment for blue whales, half-normal with sea state, group size and observer platform height as covariates for fin whales and half-normal with swell, platform height and swell as covariates for humpback whales. Selected models for all 3 species were good fits to the distance data (Craemer von-Mises test p-values 0.70, 0.31, and 0.59 respectively). Estimated density varied by season in a similar manner to the sighting rates, with strong inter-annual variability (Figure 2).

**Figure 2.**
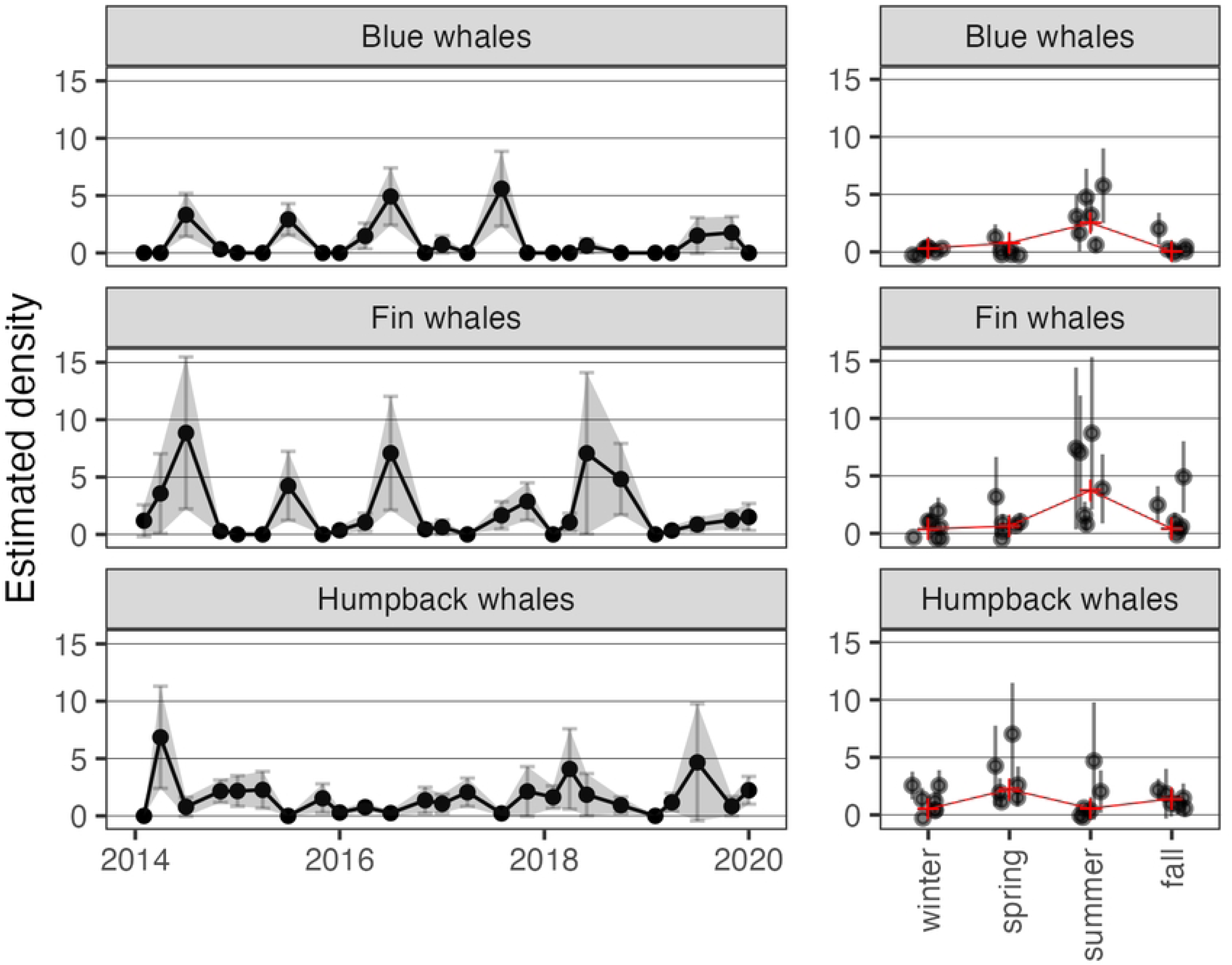
Estimated density (number of individuals per 1000 km^2^) over time across years (left) and by season (right) for each whale species; seasonal averages are shown in red.

**Table 2.** Number of whale sightings per cruise among whale species of interest.

| Year | Season | Survey start | Survey end | fin whale sightings | humpback whale sightings | blue whale sightings |
| --- | --- | --- | --- | --- | --- | --- |
| 2014 | winter | 2014-01-29 | 2014-02-04 | 2 | 0 | 0 |
|  | spring | 2014-04-03 | 2014-04-15 | 3 | 13 | 0 |
|  | summer | 2014-07-06 | 2014-07-21 | 17 | 2 | 6 |
|  | fall | 2014-11-09 | 2014-11-22 | 1 | 8 | 1 |
| 2015 | winter | 2015-01-16 | 2015-01-29 | 0 | 5 | 0 |
|  | spring | 2015-04-05 | 2015-04-18 | 0 | 4 | 0 |
|  | summer | 2015-07-08 | 2015-07-24 | 12 | 0 | 9 |
|  | fall | 2015-10-29 | 2015-11-11 | 0 | 4 | 0 |
| 2016 | winter | 2016-01-08 | 2016-01-21 | 1 | 2 | 0 |
|  | spring | 2016-04-02 | 2016-04-14 | 3 | 3 | 3 |
|  | summer | 2016-07-11 | 2016-07-25 | 3 | 1 | 10 |
|  | fall | 2016-11-07 | 2016-11-20 | 1 | 2 | 0 |
| 2017 | winter | 2017-01-06 | 2017-01-19 | 1 | 4 | 1 |
|  | spring | 2017-03-29 | 2017-04-12 | 0 | 6 | 0 |
|  | summer | 2017-08-01 | 2017-08-13 | 4 | 1 | 10 |
|  | fall | 2017-11-10 | 2017-11-23 | 3 | 4 | 0 |
| 2018 | winter | 2018-02-02 | 2018-02-09 | 0 | 3 | 0 |
|  | spring | 2018-04-06 | 2018-04-20 | 2 | 9 | 0 |
|  | summer | 2018-06-09 | 2018-06-22 | 4 | 3 | 2 |
|  | fall | 2018-10-15 | 2018-10-27 | 5 | 4 | 0 |
| 2019 | spring | 2019-04-02 | 2019-04-16 | 1 | 2 | 0 |
|  | summer | 2019-07-12 | 2019-07-25 | 3 | 3 | 4 |
|  | fall | 2019-11-05 | 2019-11-17 | 2 | 1 | 4 |
| 2020 | winter | 2020-01-05 | 2020-01-19 | 3 | 5 | 0 |

### 3.2 ESTIMATED RELATIONSHIPS

Our analysis identified highly sparse well-fitting models, selecting stable sets (Eqn. 2) comprising between 23 and 60 ASVs depending on marker and whale species that explain an estimated 81-99% of variation in density estimates after adjusting for seasonality (Table 3). Optimal model hyperparameters varied slightly; models used between 4 and 11 latent components (a maximum of 12 were available for hyperparameter optimization). The selected ASVs spanned 7-19 classes, 8-25 orders, and 9-28 families, again depending on target species and marker.

**Table 3.**
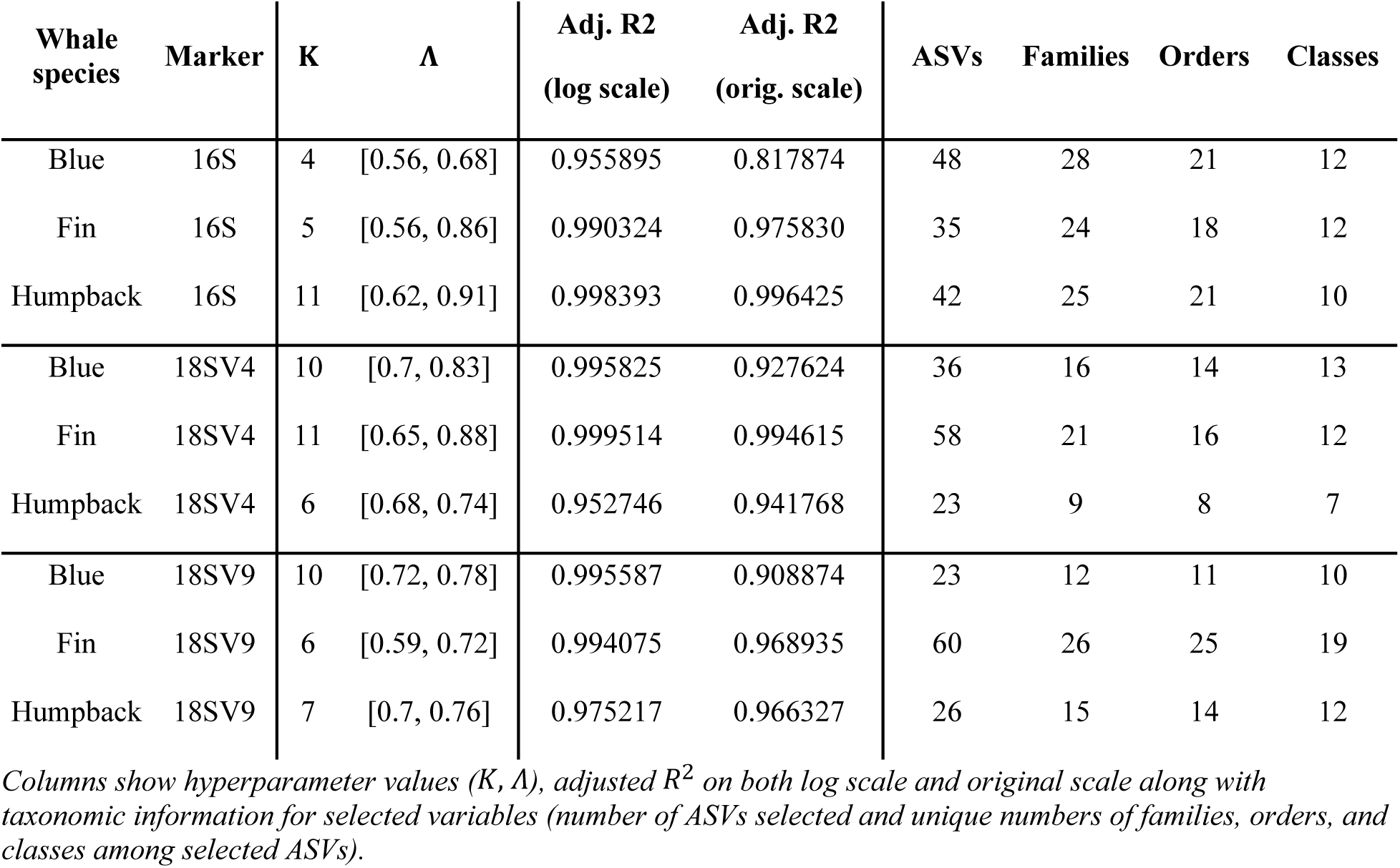
Fit summaries of all nine models selected in the analysis.

| Whale species | Marker | K | $\Lambda$ | Adj. R<sup>2</sup><br>(log scale) | Adj. R<sup>2</sup><br>(orig. scale) | ASVs | Families | Orders | Classes |
| --- | --- | --- | --- | --- | --- | --- | --- | --- | --- |
| Blue | 16S | 4 | [0.56, 0.68] | 0.955895 | 0.817874 | 48 | 28 | 21 | 12 |
| Fin | 16S | 5 | [0.56, 0.86] | 0.990324 | 0.975830 | 35 | 24 | 18 | 12 |
| Humpback | 16S | 11 | [0.62, 0.91] | 0.998393 | 0.996425 | 42 | 25 | 21 | 10 |
| Blue | 18SV4 | 10 | [0.7, 0.83] | 0.995825 | 0.927624 | 36 | 16 | 14 | 13 |
| Fin | 18SV4 | 11 | [0.65, 0.88] | 0.999514 | 0.994615 | 58 | 21 | 16 | 12 |
| Humpback | 18SV4 | 6 | [0.68, 0.74] | 0.952746 | 0.941768 | 23 | 9 | 8 | 7 |
| Blue | 18SV9 | 10 | [0.72, 0.78] | 0.995587 | 0.908874 | 23 | 12 | 11 | 10 |
| Fin | 18SV9 | 6 | [0.59, 0.72] | 0.994075 | 0.968935 | 60 | 26 | 25 | 19 |
| Humpback | 18SV9 | 7 | [0.7, 0.76] | 0.975217 | 0.966327 | 26 | 15 | 14 | 12 |

Columns show hyperparameter values ( $K$ , $\Lambda$ ), adjusted $R^2$ on both log scale and original scale along with taxonomic information for selected variables (number of ASVs selected and unique numbers of families, orders, and classes among selected ASVs).

Our model selection procedure exhibited some sensitivity to data perturbation. Depending on the model and taxonomic level, anywhere from 22-68% of selected taxa are robust to leave-one-out data perturbations. In detail, Table 4 shows the modified Jaccard index (Eqn. 4) computed for each model at the ASV, family, order, and class levels. This measure quantifies the proportion of taxa enumerated across models fit to leave-one-out data partitions that are included more often than not in the stable set.

**Table 4.** Measures of selection consistency computed at the ASV, family, class, and order level.

| Whale species | Marker | J-index<br>(ASV) | N<br>(ASV) | J-index<br>(Family) | N<br>(Family) | J-index<br>(Class) | N<br>(Class) | J-index<br>(Order) | N<br>(Order) |
| --- | --- | --- | --- | --- | --- | --- | --- | --- | --- |
| Blue | 16S | 0.353 | 331 | 0.518 | 56 | 0.682 | 22 | 0.604 | 48 |
| Fin | 16S | 0.384 | 172 | 0.488 | 41 | 0.684 | 19 | 0.514 | 35 |
| Humpback | 16S | 0.319 | 119 | 0.229 | 35 | 0.267 | 15 | 0.258 | 31 |
| Blue | 18SV4 | 0.405 | 163 | 0.563 | 32 | 0.571 | 21 | 0.556 | 27 |
| Fin | 18SV4 | 0.405 | 232 | 0.548 | 42 | 0.485 | 27 | 0.500 | 36 |
| Humpback | 18SV4 | 0.354 | 291 | 0.457 | 46 | 0.433 | 30 | 0.447 | 38 |
| Blue | 18SV9 | 0.390 | 413 | 0.558 | 86 | 0.579 | 57 | 0.587 | 75 |
| Fin | 18SV9 | 0.352 | 145 | 0.350 | 40 | 0.370 | 27 | 0.351 | 37 |
| Humpback | 18SV9 | 0.300 | 110 | 0.517 | 29 | 0.500 | 20 | 0.500 | 26 |

*For each taxonomic level, N gives the total number of unique taxa selected, and J is the proportion of those that* *were frequently selected across validation partitions. Hyperparameter settings were identical to those used in each* *of the nine models fit in the analysis.*

Supplementary Tables 3a-c enumerate selected ASVs by whale species and marker (*i.e.,* by model) along with taxonomic classifications, if known, and an estimated measure of association (*i.e.,* estimated model coefficient in Eqn. 1) quantifying multiplicative change in whale sightings per doubling of ASV relative abundance after adjusting for seasonality. For example, an estimate of 1.338 would indicate that every doubling of the relative abundance of that particular microbe (relative to its seasonal average) is associated with an estimated 33.8% increase in estimated blue whale sightings (relative to its seasonal average). Estimates greater than 1 indicate a positive association, and estimates less than 1 indicate a negative association.

### 3.3 MODEL PREDICTIONS

The selected amplicons summarized in the previous section and enumerated fully in the supplementary information (Supplementary Tables 3a-c) were strongly predictive of deviations of whale sightings from seasonal averages (Table 5). Combining predicted deviations with estimated seasonal trends, out-of-sample predictions of density were within 0.69-1.41 individuals per 1000 *km*^2^ of observed values on average, depending on whale species and marker. For context, this result represents a 11-52% reduction in prediction error compared with imputing the seasonal average, and a 20-65% reduction in prediction error compared with carrying forward the last observation from the same season, again depending on whale species and marker. While all markers produced comparable gains in predictive power, the selected 18S-V9 amplicons gave the best predictions for all target species.

**Table 5.** Summaries of predictive performance of each of the nine models selected in the analysis.

| Whale species | Marker | Correlation<br>(log scale) | RMSPE<br>(log scale) | Correlation<br>(orig. scale) | RMSPE<br>(orig. scale) | % reduction<br>(lag) | % reduction<br>(mean) |
| --- | --- | --- | --- | --- | --- | --- | --- |
| blue | 16S | 0.675 | 1.246 | 0.734 | 1.078 | 20.7 | 11.5 |
| fin | 16S | 0.939 | 0.777 | 0.907 | 1.096 | 64.7 | 52.1 |
| humpback | 16S | 0.774 | 0.838 | 0.663 | 1.238 | 42.0 | 25.8 |
| blue | 18SV4 | 0.876 | 0.839 | 0.795 | 0.990 | 27.2 | 18.7 |
| fin | 18SV4 | 0.851 | 1.025 | 0.830 | 1.415 | 54.4 | 38.2 |
| humpback | 18SV4 | 0.810 | 0.765 | 0.672 | 1.203 | 43.6 | 27.9 |
| blue | 18SV9 | 0.855 | 0.885 | 0.917 | 0.698 | 48.6 | 42.7 |
| fin | 18SV9 | 0.866 | 0.986 | 0.902 | 1.101 | 64.5 | 51.9 |
| humpback | 18SV9 | 0.876 | 0.655 | 0.746 | 1.078 | 49.4 | 35.4 |

*Correlations and root mean square prediction error (RMSPE) are reported on both the logratio scale and the original scale. Percent reductions report the relative reduction in prediction error, on the original scale, of each model compared with carry-one-forward predictions (lag method) and seasonal averages (mean method).*

Figure 3 shows leave-one-out predictions from all nine models compared with observed values. In particular, our models predicted spikes in density and low density with comparable accuracy. Predictions were made with comparable precision – as measured by 90% bootstrap percentile intervals – on the log scale, which translates to greater uncertainty associated with predictions of higher sightings.

**Figure 3.**
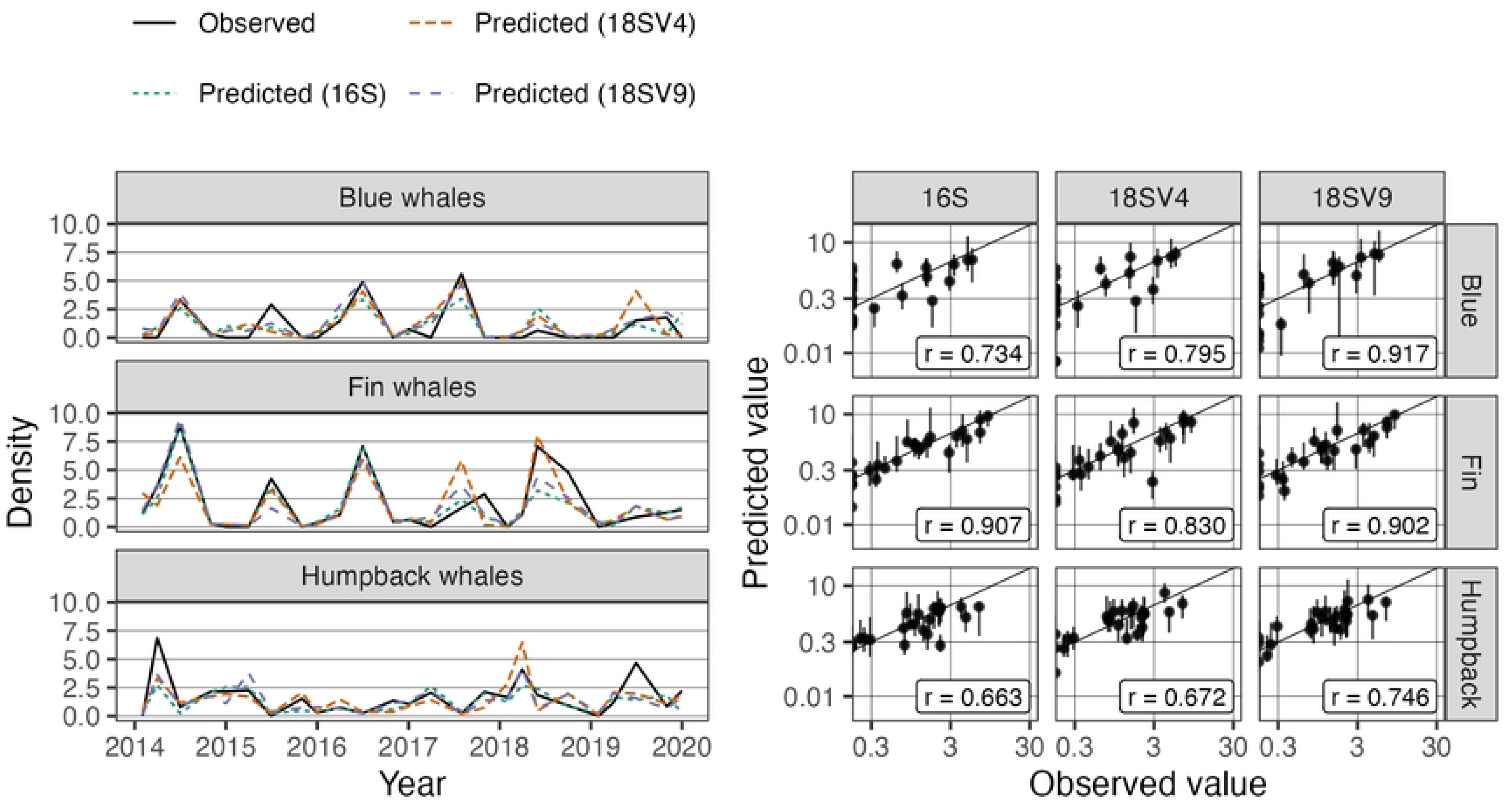
Predicted density compared with observations for each whale species and marker shown as time series (left) and simple scatterplots (right). At left, separate time series are shown for each marker and distinguished by color and line dash. At right, separate panels compare the observations and predictions for each species/marker combination on the log scale; the solid line represents perfect predictive accuracy. The vertical line ranges show 90% bootstrap percentile intervals quantifying prediction uncertainty.

### 3.4 COMMUNITIES OF TAXONOMIC ANNOTATIONS

A total of 148 unique taxonomic annotations were identified as predictive of baleen whales (Supplementary table 4b). 20% of annotations (29) were shared across all three species (blue, humpback, and fin whales), and 21% (31) of annotations were shared by two species (either blue/fin, blue/humpback, fin/humpback). Additionally, the rest (59%) were unique to a single species – 27 annotations were unique to blue whales, 43 annotations were unique to fin whales, and 18 annotations were unique to humpback whales (Supplementary table 4b).

### 3.5 NARRATIVE REVIEW FINDINGS

We found 18 publications that documented 457 microbial and plankton taxa associated with blue, fin, and humpback whales, based on studies conducted in various regions throughout the world (Supplementary table 4b). After accounting for duplicate taxa, 403 unique taxa remained. These studies illustrate taxa that are known to be associated, either internally or externally, with fin, humpback, and blue whales. They highlight various aspects of baleen whale biology and ecology, including related to fecal, digestive, respiratory, and skin microbiomes; prey composition; and feeding habitats. They also document a range of epibiotic parasitic and commensal organisms, such as barnacles and diatoms. A complete list of taxa known to be associated with whales from the narrative review is contained in Supplementary table 4b.

### 3.6 OVERLAP OF ANNOTATIONS WITH LITERATURE IDENTIFIED IN NARRATIVE REVIEW

Of the 148 unique taxonomic annotations that comprised the microbial communities predictive of baleen whales in our study, 23% of the annotations (34 out of 148) were found within the existing literature exploring baleen whale microbial parasites, commensals, prey, or respiratory-associated microbes, matching at the genus (17 out of 148) or family level (17 out of 148) (Supplementary tables 4a-b). The rest of the annotations either matched at higher taxonomic levels, including order, class, and phylum (41%; 61 out of 148) or did not have any associated matches in the literature (36%; 53 out of 148) (Supplementary tables 4a-b).

## 4 DISCUSSION

In this work we leveraged microbial and small plankton community composition for prediction of blue, fin, and humpback whale density. We found that biological communities, as identified from marker genes capturing both prokaryotic and eukaryotic plankton, were good predictors of whale density across multiple seasons and years. To our knowledge, these are the first data showing that the ecological habitat of baleen whales can predict their density and track interannual variability. These results align with a growing body of evidence suggesting that baleen whale distributions in the California Current are tightly coupled to environmental conditions that drive prey availability (Palacios et al. 2019, Irvine et al. 2025, Ryan et al. 2025), which in turn are linked to microbial and planktonic community composition.

From among six to eight thousand candidate microbial taxa per genetic marker (Supplementary Tables 2a-c), our analyses found small groups of taxa that represent the ecological habitat of baleen whales and are strongly associated with and strongly predictive of the density of blue, fin, and humpback whales (Table 3, Table 5, and Supplementary tables 3a-3c). In particular, we identified groups of 23-60 distinct ASVs per whale species that explain 81-99% of nonseasonal variability in estimated whale density (Table 3) and predict density estimates to within ∼1 individual per 1000 *km^2^* (Table 5). The predictive accuracy achieved by our approach improves out-of-sample RMSE by up to an estimated 65% compared with naive methods (Table 5).

In total, these groups of ASVs represented 148 unique taxonomic annotations identified across marker genes, with 20% shared among all three whale species, 21% shared between two species, and 59% unique to a single species. These results suggest that there is some overlap in the microbial ecological habitat among blue, humpback, and fin whales, but each species is also related to a distinct microbial community, with blue whales exhibiting the highest uniqueness in microbial taxa.

Additionally, we compared the predictive microbial annotations from our study to taxa found in our review of previously documented in the literature on known microbe and baleen whale connections. We found that 23% of the predictive taxa matched taxa from the literature we reviewed, at the genus or family level. Many of the taxa that matched are known to be prey, parasites, and commensals of baleen whales. For example, members of the genera *Sphingomonas*, *Pseudoalteromonas*, *Acinetobacter*, and *Pseudomonas* have also been found in blow samples of humpback whales (Apprill *et al*. 2017). Additionally, members of the genus *Psychrobacter* have also been found on skin and in blow samples of humpback whales (Apprill *et al*. 2014, Apprill *et al*. 2017) and in blow samples of blue whales (Dominguez-Sanchez *et al*. 2022). *Serratia* spp. have also been observed in the digestive tract of fin whales (Herwig & Staley 1986). Additionally, both calanoid and cyclopoid (including Poecilostomatoida) copepods have been detected as prey in humpback whale feces (Reidy *et al*. 2022). In addition to these examples, many other taxa in our communities were also found in the existing literature on microbial whale associates (Supplementary tables 4a-b). These findings suggest that some of these microbial signals may be ecologically relevant—either directly through food-web or symbiotic interactions, or indirectly by reflecting environmental features of whale habitats, such as distinct water masses or seasonal patterns (Supplementary tables 4a-b). Given that the rest either matched at higher taxonomic levels (41%) or had no documented matches in our literature search (36%), and some were uncultured or unassigned, this suggests that future work could examine these specific taxa further.

Ultimately, our focus is not on individual ASVs in isolation, but rather on their combined presence as a collective community or consortium. This suite of microbes and small plankton likely represents an integral component of the ecological habitat surrounding baleen whales, contributing to the complex collection of organisms that may influence or reflect their density. Our results point to a consistent community-level association between large cetaceans and microbes or small plankton, which may reflect physical and chemical signatures of oceanic water masses (Agogué *et al*. 2011, Samo *et al*. 2012, Djurhuus *et al*. 2017, Saunders *et al*. 2022). Marine microbial diversity is vast and poorly characterized at the strain level; thus, we refrain from presenting individual microbial taxa as predictors of whales. However, microbial consortia are well-known to transform nutrients and other small molecules in ways that shape the surrounding food web (Gralka *et al*. 2020, Kieft *et al*. 2021, Dal Bello *et al*. 2021). We therefore postulate that there exists a microbial community index that is characteristic of habitat favorable to whale species, whether due to indirect relationships (e.g., because it reflects the biochemical habitat of prey items like krill and fish) or some direct relationships (e.g., microbial associates of whales).

Notably, our molecular data encompass both prokaryotic and eukaryotic plankton, which allowed us to characterize microbial diversity across different domains of life. There are likely many bacterial-bacterial and bacterial-plankton interactions, as well as other as-yet uncharacterized ecological relationships among the predictor taxa. Taken together, the community-level synergy is likely greater than the sum of its parts, and we present a first step at understanding and untangling the complex linkages between the food web base and its apex, as well as other ecological relationships.

Our statistical analysis framework combines logratio methodology for compositional data analysis (Aitchison & Bacon-Shone 1984) with sparse partial least squares (Chun & Keleş 2010) for interpretable dimension reduction, and we use stability selection (Meinshausen & Bühlmann 2010) to improve the robustness of the variable selection procedure to small perturbations in training data. We found that the resulting estimation procedure in our analysis exhibits a range of selection consistencies, depending on the model. Among the 9 models in our analysis, anywhere from an estimated 30-40% of ASVs were consistently selected when holding out one data point at a time (Table 3). Although surprisingly low, given that stability methods aim precisely at achieving consistent selection, there are several possible explanations for this result. First, the modest sample size entails that each data point exerts considerable influence on model fit. Among the 25 cruises in our analysis, one observation constitutes 4% of available data. Second, the density estimates are sparse – most cruises record few sightings and the limited number of cruises per year (∼4 per year) may mean that we miss the full range of whale densities in a given year (e.g., minimum or maximum). Although spikes in sightings comprise only 8-16% of available data, depending on whale species, it is reasonable to speculate that these cruises capture the most information about potential ecological correlations. By comparison with the remaining 84-92% of the data, removal of one or two of these high-sightings observations substantially alters the variation in the time series. Thus, depending on which observations are held out, fitted models may describe fundamentally different ecological processes – either small variations among low density or large fluctuations between high and low scaled sightings. Third, our analysis did not account for (a) uncertainty in taxonomic classification of ASVs or (b) potential biases inherent to eDNA methods. These include extraction, primer, and amplification biases, which can cause some species to be missed and others to appear more or less abundant than they really are. Strong correlations among amplicon relative abundances due to either factor could lead to instability in variable selection under small perturbations of the data; this is a well-documented phenomenon in the statistical literature. Fourth, partial least squares estimation is known to be sensitive to outliers -a fact which has produced proposals for robust PLS estimators (Hoffman et al. 2015). It is therefore plausible that sparse partial least squares would exhibit high selection variability under data perturbations, especially in light of the data sparsity discussed above. Finally, varying uncertainty in density estimation by cruise (Figure 2) is not accounted for in the modeling framework, but may produce uneven signal strength of associations between microbial communities and baleen whale density depending on which cruises are used to fit models. All of these factors may contribute to the wide range of selection consistency observed in our work.

Overall, our findings suggest that microbial communities can help to predict baleen whale densities and that some microbial community predictors may be ecologically relevant. However, further research is needed to more holistically examine the groups of ASVs that collectively define these microbial communities. This could also include investigating some of the taxa identified in this study in greater detail to clarify their functional roles and identify ecological relationships with baleen whales, applying other genetic methods to improve taxonomic resolution, or further exploring novel or understudied microbial associations relevant to whale habitats. Using microbial community composition to characterize the potential ecological habitat of baleen whales can enhance predictive models and inform hypotheses about the ecological relationships between whales and bacterioplankton, phytoplankton, and zooplankton. We believe that this approach is broadly transferable to other ecological systems where microbial data are collected in parallel with observations of larger organisms and may provide insight into ecological relationships and potential of the ecological habitat to predict their distribution and abundance.

## ACKNOWLEDGMENTS

We are grateful to Matthew Robbins for his support in developing code to programmatically retrieve taxonomic information from the World Register of Marine Species (WoRMS) about taxa in our literature review. This research is partly based upon work supported by the Research, Scholarly & Creative Activities Program awarded by the Cal Poly Division of Research. EVS was supported by a partnership among CalCOFI participants, including Scripps Institution of Oceanography (SIO), NOAA Southwest Fisheries Science Center (SWFSC), California Department of Fish and Wildlife, and California Sea Grant.

## AUTHOR CONTRIBUTIONS

**Erin V. Satterthwaite**: conceptualization; methodology; writing – original draft; writing – review and editing; project administration. **Trevor D. Ruiz**: formal analysis; methodology; validation; visualization; software; writing – original draft; writing – review and editing; project administration. **Nastassia V. Patin**: conceptualization; writing – original draft; writing – review and editing. **Michaela N. Alksne**: conceptualization; methodology; data curation; writing – original draft; writing – review and editing. **Len Thomas**: methodology; formal analysis; software, writing – original draft; writing – review and editing. **Julie Dinasquet**: conceptualization; writing – review and editing. **Robert H. Lampe**: investigation; data curation; writing – original draft; writing – review and editing. **Katherine G. Chan**: methodology; formal analysis; software. **Nicholas A. Patrick**: methodology; formal analysis; visualization; software. **Andrew E. Allen**: conceptualization; investigation. **Simone Baumann-Pickering**: investigation; writing – review and editing. **Brice X. Semmens**: conceptualization; writing – review and editing.

## DATA AVAILABILITY STATEMENT

Sequence data generated in this study have been deposited in the NCBI Sequence Read Archive under BioProject accession numbers PRJNA555783, PRJNA665326, and PRJNA804265. Sighting data, density estimates, NCOG sample metadata, taxonomic annotations, and data used for modeling are publicly available via Zenodo at https://doi.org/10.5281/zenodo.15678927. R code to reproduce the analyses is available on GitHub at https://github.com/ruizt/marine-mammal-edna.

## SUPPLEMENTARY MATERIALS

### SUPPLEMENTARY TABLES

Supplementary Table 1: numbers of NCOG samples tabulated by cruise and transect.

Supplementary Table 2a-c: taxonomic annotations of candidate ASVs used in the analysis for (a) 16S, (b) 18SV4, and (c) 18Sv9 markers.

Supplementary Tables 3a-c: estimated model coefficients and taxonomic annotations of selected ASVs used for prediction for (a) 16S, (b) 18SV4, and (c) 18Sv9 models.

Supplementary Tables 4a-b: known direct relationships between blue, humpback, and fin whales and bacteria, microbes, and small plankton from existing literature; comparison with potential relationships identified by our models.

Supplementary Table 5a-b: (a) per-species sample size of detections before and after truncation for line transect analysis; (b) list of all line transect detection function models fitted for each species, with corresponding AIC values and delta AIC (i.e., the difference in AIC between that model and the lowest-AIC model for that species)..

### SUPPLEMENTARY FIGURES

Supplementary Figure 1: residual diagnostic checks for density models.

